# Global DNA Hypomethylation as a Biomarker of Accelerated Epigenetic Ageing in Primates

**DOI:** 10.1101/2024.03.27.586982

**Authors:** Michael T.S. Girling, Nofre M Sanchez, Ursula M. Paredes

## Abstract

**Introduction:** Epigenetic clocks based on DNA methylation patterns provide a powerful tool for measuring biological ageing, but requiring genome-wide methylation data and high costs limits their broad application across species and populations.

**Methods:** We investigated whether simply quantifying global DNA methylation levels could serve as an inexpensive proxy for epigenetic ageing, using a captive colony of owl monkeys (*Aotus nancymaae*) using a colorimetric ELISA assay to measure proportional content of levels of blood and brain 5-methylcytosine (5-mC) across the genome, comparing owl monkeys with known exposures to ageing accelerators and controls.

**Results:** we found that global 5-mC declined significantly with chronological age in blood, and in the brain of parents. Notably, this age-related blood hypomethylation in individuals experiencing early life maternal rejection was accelerated. Parenting experience also accelerated DNA methylation loss with age, but this effect was specific to the brain and not seen in blood. Infection history did not impact blood 5-mC trajectories. Although multiple regression models did not replicate all findings, likely due to sample size constraints, our results demonstrate that global DNA hypomethylation tracks biological ageing in blood.

**Discussion:** This simple metric successfully detected accelerated epigenetic ageing induced by early adversity, as well as distinct patterns relating to reproductive investment in the brain - phenotypes typically identified by sophisticated epigenetic clocks. Quantifying global methylation thus provides a cost-effective alternative approach to assessing susceptibility to environmentally-driven accelerated ageing across primate species and populations where DNA methylation arrays or sequencing are impractical.

## 1. Introduction

Predicting poor health outcomes in primate populations is of interest for those involved in the conservation and preservation of primate welfare. One predictor of diminishing health and increased mortality across primates is increased chronological age (Bronikowski et al., 2011). Alteration and loss of epigenetic information in a genome is one well studied cause of mammalian ageing (Fraga & Steller 2007; Yang et al., 2023) and it is associated with increased mortality (Marioni et al., 2015). In the last decade studies have shown that changes in DNA methylation levels in a group of genomic loci predict an individual’s age (Hannum et al., 2013; Horvath and Raj 2018, Bell et al., 2019). Not all age equally, those exposed to harsh environments and poor resource availability age faster (reviewed in Kenyon, 2010, and in Pepper et al., 2018). Accordingly, epigenomic clocks become ‘accelerated’ in response to stress exposure (McCrory et al., 2022), pathogens (Horvath & Levine 2015), lifestyle habits (Gao et al., 2016) and toxins (Xiao et al., 2021), dietary changes, psychiatric illness, amongst others in humans (reviewed in Oblak et al, 2021). Then application of genomic tools could help identify wild populations at increasing chances of suffering from environmentally triggered frailty and disease (Theissinger et al., 2023). Pan primate epigenomic clocks have been created (Horvath et al., 2020) which could answer this need. However, there is little overlap between the availability of financial and bioinformatic resources and expertise needed to apply epigenetic clocks and primate habitat nations primarily in the developing south (Davila et al., 2004) where conservation of animals is applied and where increasingly, primate populations are declining (Estrada et al., 2017). Low-cost, and low-tech alternative solutions are needed to bridge this gap.

Total 5-methylcytosine (5-mC) content relative to total cytosine across the genome also decreases with age (Wilson & Jones 1983, Calvanese et al., 2009). Global DNA methylation content measured using an ELISA, can be measured on any primate genome, irrespective of availability of a fully sequenced genome or genome biologists, and can be carried out in basic diagnostics laboratories. Furthermore, global DNA methylation loss is also a hallmark of tumours (De Smet and Loriot 2010), is associated with poor cancer prognosis (Li et al., 2014), and increased risk for some immune and psychiatric illness (reviewed by Pogribny & Beland 2009). Global loss of DNA methylation has also been documented in populations exposed to environmental disruptors and stressors (in humans Glad, et al., 2017) vertebrates (wall lizards Paredes et al., 2016; alligators Nilsen et al., 2016, hyenas Laubach et al., 2021) and specifically primates (squirrel monkeys, Seenayah et al., 2024). This evidence, together, demonstrates significant parallels between ageing and responses between refined epigenomic clocks and simple global DNA methylation. Measurement of global DNA methylation might also offer insight into accelerated ageing.

An important question is since, cognitively and socially complexity is a salient trait in primates, stressful environments might negatively affect the sensitive brains of primates. As tissues age by accumulating damage with time, and cell types display different lifespans (Kline & Cliffton 1952, Nowakowski, 2006), cellular functions, vulnerabilities (Horvath et al., 2022) and evolutionary constraints, which might impact how they age, it is likely that blood ageing and responses differ from the brain responses to the same exposures. Some evidence supports this. Epigenomic ageing clocks which allow prediction of ageing across tissue types exist (Horvath et al., 2013). However, brain cortex specific epigenetic clocks have been developed (Shireby et al., 2020) which demonstrate the differences between blood and cortical brain ageing in humans (Grodstein et al., 2021). Outside clocks, there is evidence that for some primate brains, epigenomes might age more slowly than blood (in chimpanzees and humans, Guevara et al., 2022).

We propose that measuring global DNA methylation could help identify primate undergoing accelerated epigenetic ageing, and what’s more, distinguish tissue-specific ageing vulnerabilities to ageing exposures. We tested these hypotheses in an ageing population of owl monkeys (*Aotus nancymaae*) from the IVITA captive breeding colony in Peru, using whole blood and postmortem brain samples. This colony was studied as it has individual records on health, environmental exposures and reproductive records spanning 40 years allows us to interrogate interaction ageing trajectories of global DNA methylation and life exposures. Since data comes from a managed colony, factors such as differences in nutrition or environmental fluctuations, known contributors to DNA methylation fluctuations are kept constant.

In this paper we compared measures of global DNA methylation of whole blood and postmortem brain of owl monkeys with and without known history of exposure to epigenomic clocks accelerators: early life adversity (Chaudhari et al., 2022) infections (Horvath & Levine, 2015) and parity investment (Ryan et al., 2018). We predict that when facing the same ageing exposures global DNA methylation age related decline between blood and brain samples would differ, and that age related decline would be accelerated by exposure to above described exposures.

## 2. Methods

### a. Samples/Materials

This study used IVITA Owl monkeys (*Aotus nancymaae*) samples from the IVITA colony, from San Marcos University (UNMSM) in Iquitos, Peru. The materials utilised in this study comprised Owl monkey population housed in the Centre for conservation and reproduction of primates of the veterinary Institute of tropics and andes (IVITA from their acronym in spanish) from San Marcos University (UNMSM), located in the city of Iquitos, Peru. Whole blood samples were also obtained using standard phlebotomy techniques, obtained during standard sanitary controls for the colony where 2cm diameter drops were stored in filter paper (Whatman® 903 Protein saver card) and stored in the freezer at -20oC until processing. A total of 48 blood samples were included, with age distribution mean=10.02 years, ranging between 3 and 27 years (SD=5.40), 23 females, 25 males were included.

Brain samples of the left frontal cortex are obtained from individuals who died from natural causes, 0-2 hours post-mortem window, which were stored in 100% ethanol until the moment of extraction. The total sample was n=41, with 16=males, 17=females, mean age= 9.46 years, ranging 2 and 31 years (SD=8.20)

All DNA extraction prior to this project was conducted with DNeasy blood and tissue kit (Qiagen, UK). DNA quality and quantity were quantified using a Nanodrop spectrophotometer (2.0) in the Laboratory for Ecology and Biodiversity in Cayetano Heredia University in Lima, Peru.

### b. Quantification of global DNA methylation using ELISA

The MethylFlash Global DNA Methylation (5-mC) ELISA Easy Kit (Epigentek, USA), utilising a colourimetric assay, was employed for brain and blood samples. The ELISA assays were conducted at the Blizard Research Institute of Queen Mary University. Ethical approval of procedures was previously awarded to Dr. Paredes for the conduction of this work at QMUL. Ethical permits were obtained from the Peruvian authority by UNMSM IVITA staff to use these samples for research purposes (AUT-IFS-2021-040 (SERFOR RDG NO D000334-2021-MIDAGRI-SERFOR-DGGSPFFS)).

Raw optic density measures were obtained by measuring colour intensity produced during ELISA using a standard plate reader at 450 nm. ODs were standardised using a standard curve using a 4-factor polynomial curve to obtain a slope. This slope was then used to convert absolute OD values into absolute global DNAm percentages.

### c. Data/Statistical analysis

A linear regression analysis was conducted separately for blood and brain samples to examine the relationship between global DNAm and age while adjusting for the potential confounding effect of sex. Pearson’s correlation, quantile normalisation and visualisation of the data were performed using the base R ‘stats’ package, preprocessCore, and visualisation packages ggplot, ggpubr, cowplot and tidyr, respectively. To investigate the effect of stressors and energy investment (maternal rejection, infections and parity) on global DNAmethylation, individuals were categorised. For parity we created two categories: Individuals with parity history and no parity history. To study effects of early life adversity individuals were divided into; maternally rejected and not rejected. To investigate the effect of infections, we extracted history of treatments based on individual medical records, individuals were categorised as healthy or treated (mostly medication to treat wide spectrum antiparasitic and antibacterial infections (Ivermectin) and diagnosis for tuberculosis (tuberculin test).

All statistical computations were conducted using the statistical program R (version 4.3.0; http://www.r-project.org). All P values reported are two-tailed and statistical significance was defined as P<0.05.

## 3. Results

This study examined global DNA methylation patterns in blood and brain tissue across the lifespan of owl monkeys to understand how early life adversity, investment in reproduction and infections impact epigenetic ageing in blood and when possible, in brain tissue. Our studies show that when considering the entire cohorts, blood DNA methylation levels decreased with age, whereas brain DNA methylation levels did not. Exposures such as experiencing early life adversity accelerated the blood DNA methylation decline with age while having live offspring was associated with lower DNA methylation levels in brain samples, but not in blood. Infections were found to be not associated with differences or changes in global DNA methylation content in blood.

### a. Ageing has a significant hypomethylation effect on blood but not in brain DNA

The effect of ageing on global DNAmethylation trajectories was investigated in both tissues. Pearson’s correlation coefficient analyses indicated a statistically significant relationship between age and blood 5-mC%, although significant correlation was weak (p < 0.05, R^2^ = 0.1). The negative correlation suggests that as age increased, DNAm levels also tended to decrease. A simple linear regression analysis revealed a significant relationship between age and blood global DNAm levels (F(1,46)=5.14, p < 0.05). The negative β1 coefficient indicated that for each year of increase in age, blood DNAm levels decreased by approximately 0.33% (see Figure 1a). This correlation coefficient indicates that chronological ageing influences blood DNA methylation trajectories, however, factors other than ageing not captured by our model played a greater contribution to explain DNA methylation content.

**Figure 1.**
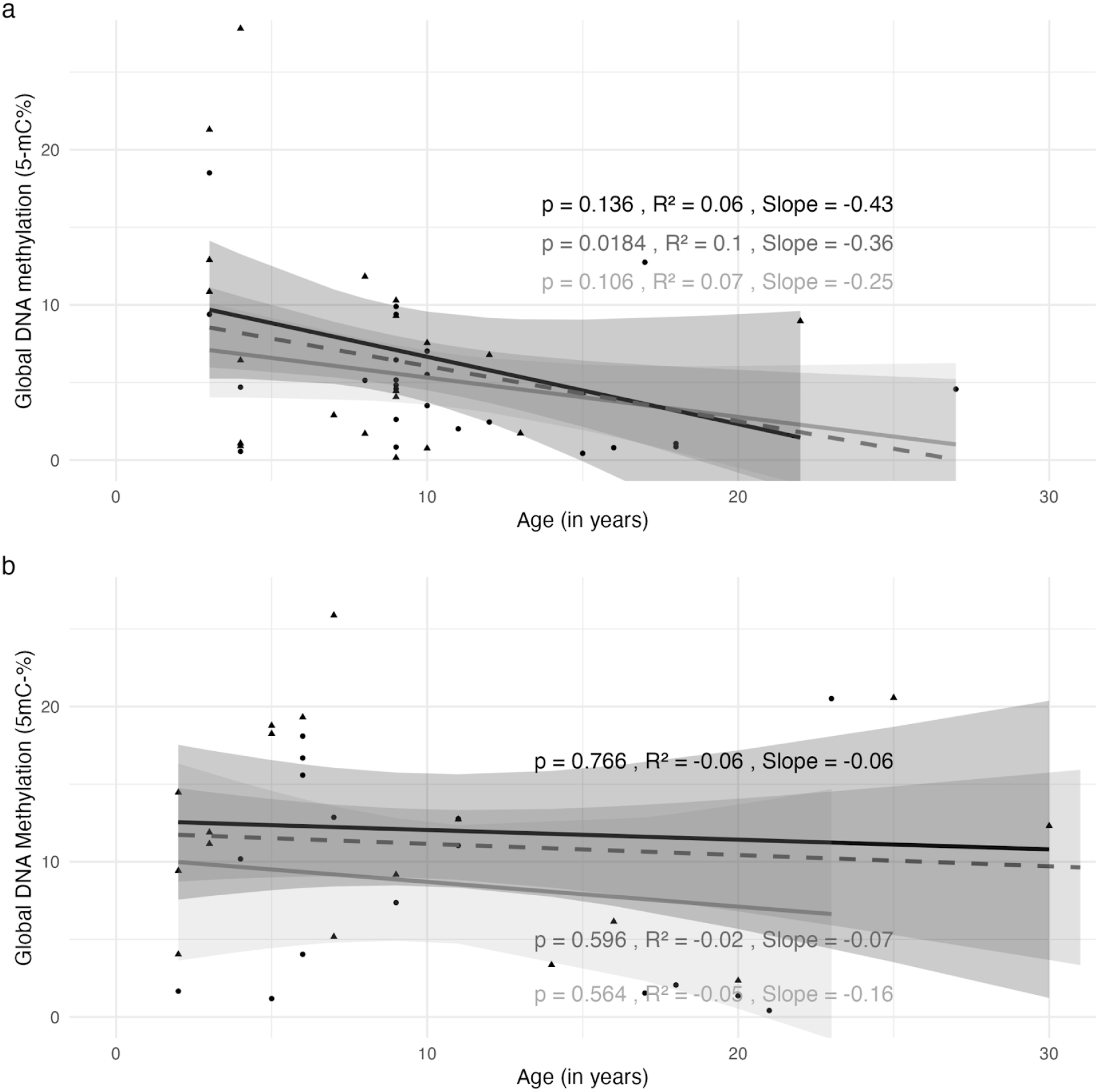
Ageing trajectories of blood and brain global DNA methylation (5-mC%) in owl monkeys. a) Scatterplot of blood global DNAm (5-mC) across owl monkeys of different ages. Males (n=25; p = 0.11) and females (n=23; p = 0.14) 5-mC%∼ age trajectory is depicted in grey and black regression lines respectively. Both combined = dashed dark grey. b) Scatterplot of brain global DNAm (5-mC%) across owl monkeys of different ages. Males (n=16; p=0.56) and females (n=17; p = 0.76) 5-mC%∼ age trajectory is depicted in grey and black regression lines respectively. Both combined = dashed dark grey. Males are represented by circle data points and females are represented by triangle data points. Linear regression analysis identified a significant and negative correlation between blood 5-mC and age (p < 0.05) and no correlation between brain 5-mC decline and increasing age (p = 0.60).

A similar comparison was conducted for brain DNA. A Pearson’s correlation coefficient analysis indicated a non-significant relationship between age and brain 5-mC% (p = 0.60, R^2^ = 0.007). The negative correlation suggests that as age increased, DNAm levels also tended to decrease. A simple linear regression analysis revealed a non-significant relationship between age and DNAm levels (F(1,39)=0.28, p = 0.60). The results indicate that in our study, brain global DNAmethylation does not significantly decline with age.

Pearson’s correlations examined sex differences in DNAm levels in both blood and brain samples. Blood 5-mC% patterns in ageing cohort did not differ between sexes, with a non significant association between age and DNAm levels in males (r = -0.31, p = 0.14), supported by a non-significant linear regression (F(1,23) = 2.37, p = 0.14). Similarly, in females we identified a nonsignificant pattern (correlation coefficient r = -0.30, p = 0.16) and a non-significant linear regression (F(1,22) = 2.13, p = 0.16) (Figure 1a).

Brain DNAm ageing patterns did not differ between sexes either, with non-significant correlations and linear regressions for both males (r = -0.26, p = 0.34; F(1,13) = 0.97, p = 0.34) and females (r = -0.11, p = 0.71; F(1,11) = 0.14, p = 0.71), shown in Figure 1b below.

### b. Brain samples has higher global DNA methylation content than blood samples

An independent two-sample t-test was performed to examine the differences in global DNAm levels between blood and brain tissues. The analysis revealed a statistically significant difference in global DNAm levels between blood and brain (5-mC_blood_=6.05, SD=5.66; 5-mC_brain_=11.20, SD=6.93; t(77.18)=-3.79, p < 0.001), showing that in owl monkeys there are differences in content of DNAm between these two tissue types that persist through the life course. As DNA methylation is affected by ageing, and blood and brain studies were not conducted in the same individuals, we controlled for age by using a linear regression. This test confirmed a significant difference between blood and brain tissues (β=5.13, 95% CI=[2.51,7.76], p < 0.001); Figure 2. The owl monkey brain samples exhibited, on average, approximately 5.13% higher global DNAm compared with blood samples, after accounting for age.

**Figure 2.**
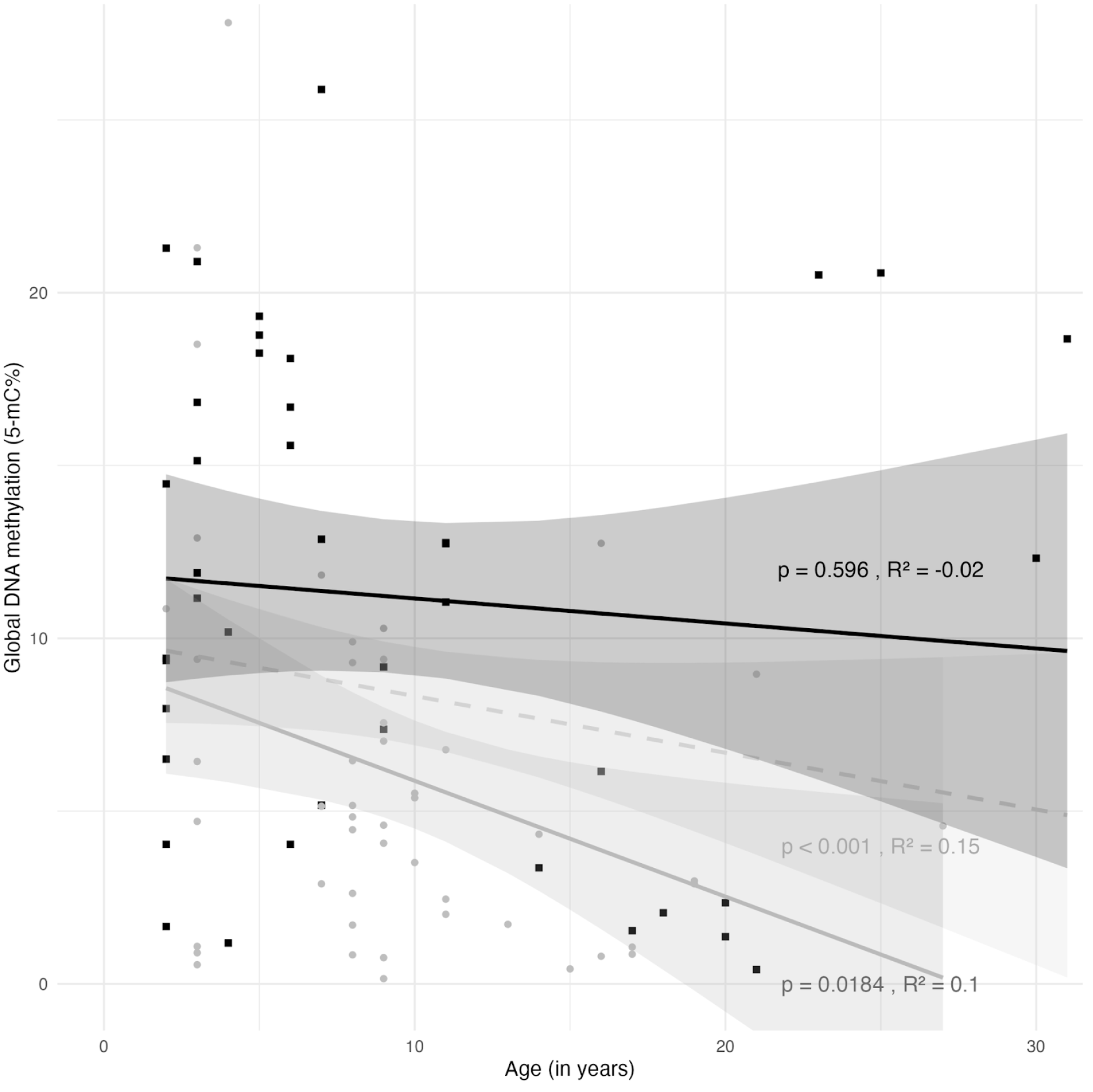
Differences in global DNA methylation (5-mC%) content in brain and blood of owl monkeys. Scatter plot showing blood and brain normalised 5-mC values (*N*=89) in an ageing cohort. Grey circles indicate blood samples, (*N*=48) and black squares indicate brain samples, (*N*=41). The grey, black, and light grey dashed lines represent the linear regression fit for blood samples, brain samples and the combined dataset, respectively. The R-squared values are -0.02, 0.1, and 0.15, respectively, indicating the proportion of variation in global DNA methylation that can be explained by age.

#### 2. Early life adversity accelerates blood global DNA hypomethylation

The IVITA colony of Owl monkeys has reported an increase in events of spontaneous maternal rejection soon after birth (Sanchez et al., 2006; Osman et al., 2020). We predicted that blood epigenetic ageing would be accelerated by this exposure appearing as steeper global DNA methylation declines with age. To test this we first compared mean 5-mC% in rejected and non rejected. We found that rejected and non rejected monkeys had similar global DNA methylation content (Rejected_5mC_ mean = 6.57, SD = 6.24, IQR = 6.40; Non rejected_5mC_ mean = 5.48, SD = 5.02, IQR = 7.27, Figure 3a). However, controlling for age, a linear regression analysis showed that both rejected and non rejected 5-mC% decline with age, but decline was only significant for the rejected (β=-0.46, 95% CI=[-0.86, -0.05], p < 0.05).

#### 3. History of Infections treatment has no effect on DNA methylation ageing patterns in blood

As repeated infections can lead to accelerated epigenomic ageing of blood tissues, we predicted that repeated infections would accelerate age-related decline of global DNA methylation in owl monkeys. Comparative analysis between two groups: individuals with a history of medical treatment for infections and those with an absence of medical histories was conducted. An independent Two Sample T-test revealed a statistically non-significant difference in global DNAm levels between healthy and previously afflicted (No infection_5-mC%_=6.46, SD=6.06) (Infection_5-mC%_=4.26, SD=2.99, t(25.56)=1.59, p = 0.16), shown in Figure 4a. We followed up with a linear regression to investigate differences in global DNAm decline with age between health and infection. This analysis showed that age had a marginally-significant negative association with global DNAm levels in both healthy individuals (β=-0.33, 95% CI=[-0.66,<0.01], p = 0.05); and a no association in those treated for infections (β=-0.49, 95% CI=[-1.31,0.32], p = 0.20), shown in Figure 4b. However, the regression line shown for individuals with infection history has a steeper slope than healthy individuals.

#### 4. Reproductive investment is associated with lower global DNA in the brain but not in blood

As investing in having offspring can impact cognitive performance in *Aotus* spp, and it has been associated with accelerated epigenomic ageing in humans we predicted that parenting experience would accelerate ageing of blood and brain. Furthermore, as blood cells and brain cells have different longevity and function, we predicted different ageing trajectories and responses to parenting between tissues. In this analysis, we only included individuals age 3 or older, which is the age at which sexual maturity is reached in owl monkeys.

First, we compared blood and brain DNA methylation between ‘with offspring’ versus ‘no offspring’, including both sexes (Figure 5a). An independent Two Sample T-test revealed non-significant differences for brain and in blood (Parents_5-mC%_=3.60, SD=3.04 Non parents_5-mC%_ =5.44, SD=3.65, t(18.93.)=-1.38, p = 0.18) but significant differences in content of DNA methylation for the brain Parents_5-mC%_= 7.41, SD=7.88; Non parents_5-mC%_=13.23%, SD=5.91, (t(16.60)=-2.19, p < 0.05).

**Figure 3.**
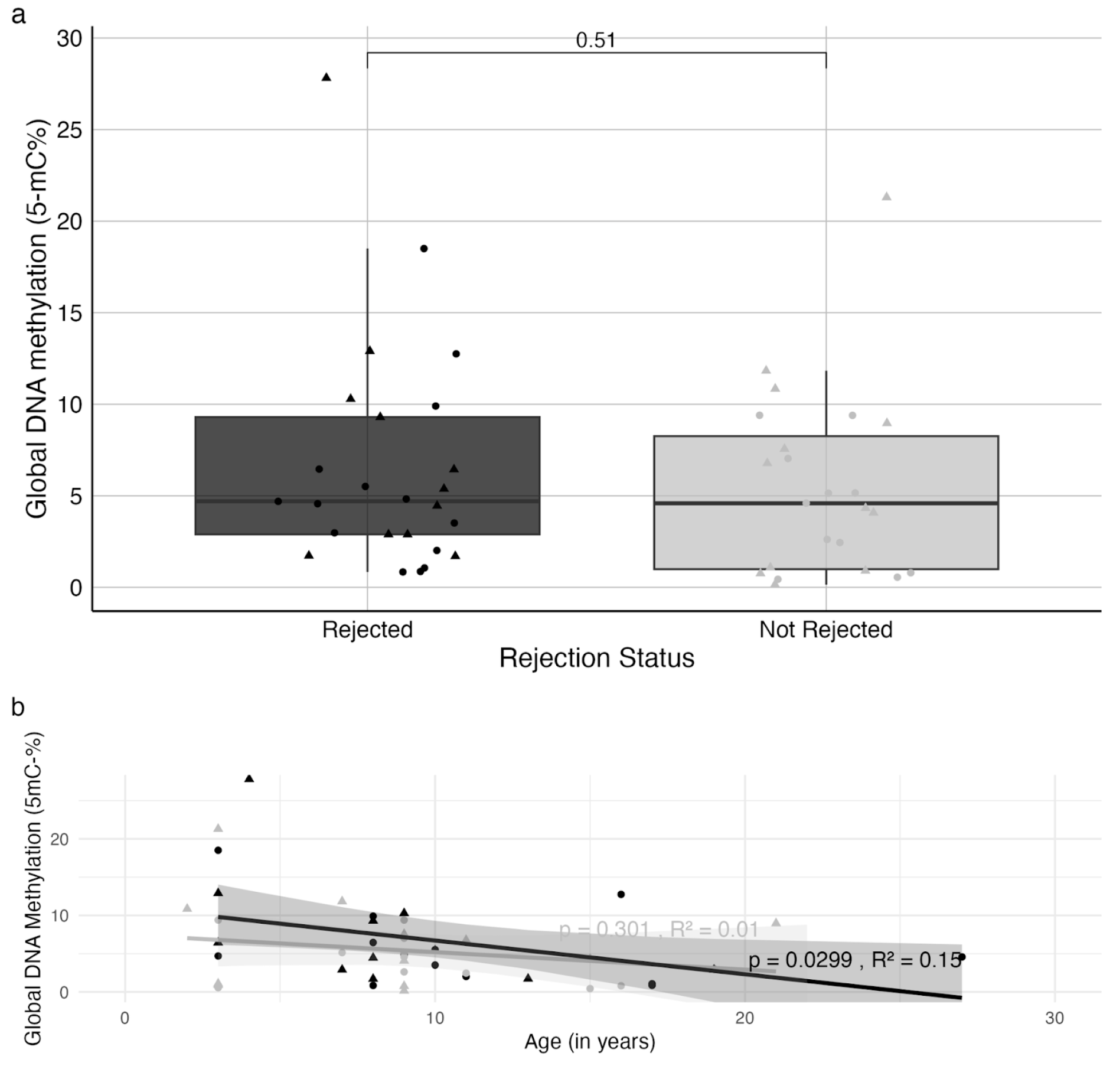
Effect of Early life adversity on global DNA methylation content and ageing trajectories. a) Boxplots showing the difference in global DNA methylation levels of blood samples between rejected (black boxplot; mean = 6.57, median = 4.70), and not rejected (grey boxplot; mean = 5.48, median = 4.59). Welch Two Sample/Student’s *T*-test show non-significant differences in overall 5-mC% (t(45.23) = -0.6675, p = 0.51). For each box, the central mark represents the median; the lines delineate the 95% confidence interval of the median; edges indicate the 25th and 75th percentiles. b) Scatterplot showing the relationship between global DNA methylation ∼ age in rejected (black regression line) and not rejected (grey regression line) owl monkey blood samples. Females and males are represented by triangles and circles respectively. Linear regressions demonstrate significant and negative relationship between 5-mC% and age in rejected but not in not rejected.

**Figure 4.**
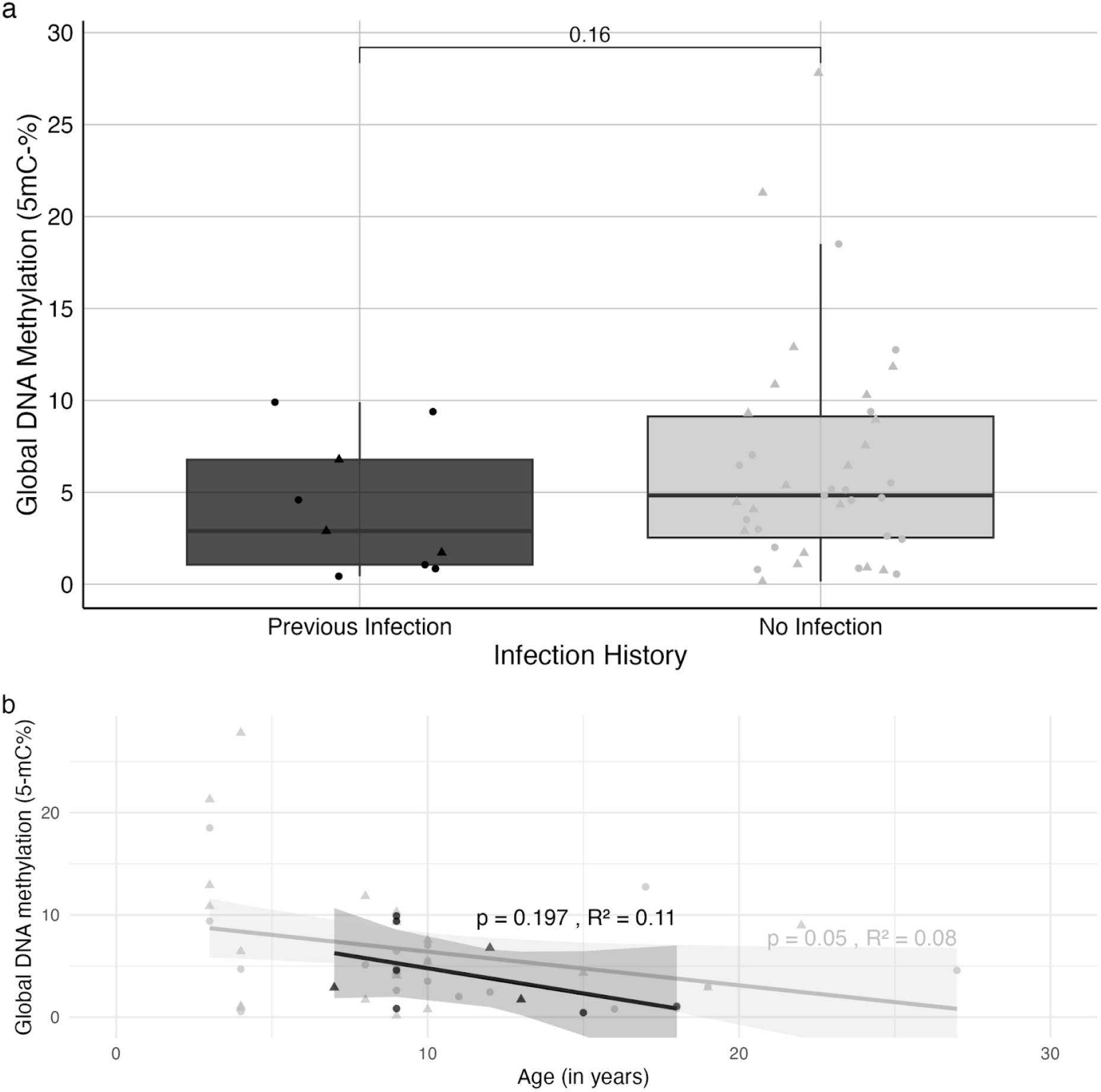
Relationship between global DNA methylation and infection history in owl monkey blood samples. a. Boxplot representing median 5-mC% individuals with and without history of infection treatment (‘previous infection’ *n*=9; ‘no infection’ *n*=39). The black boxplot represents infection; the grey box plot represents no infection. For each box, the central mark represents the median; the lines delineate the 95% confidence interval of the median; edges indicate the 25th and 75th percentiles. b) Scatterplot showing global DNAm (5-mC) interaction with age in individuals treated for infections (black regression line, *n*=9) and with no infection (grey regression line; *n*=39). Males are represented by circle data points and females are represented by triangle data points.

**Figure 5.**
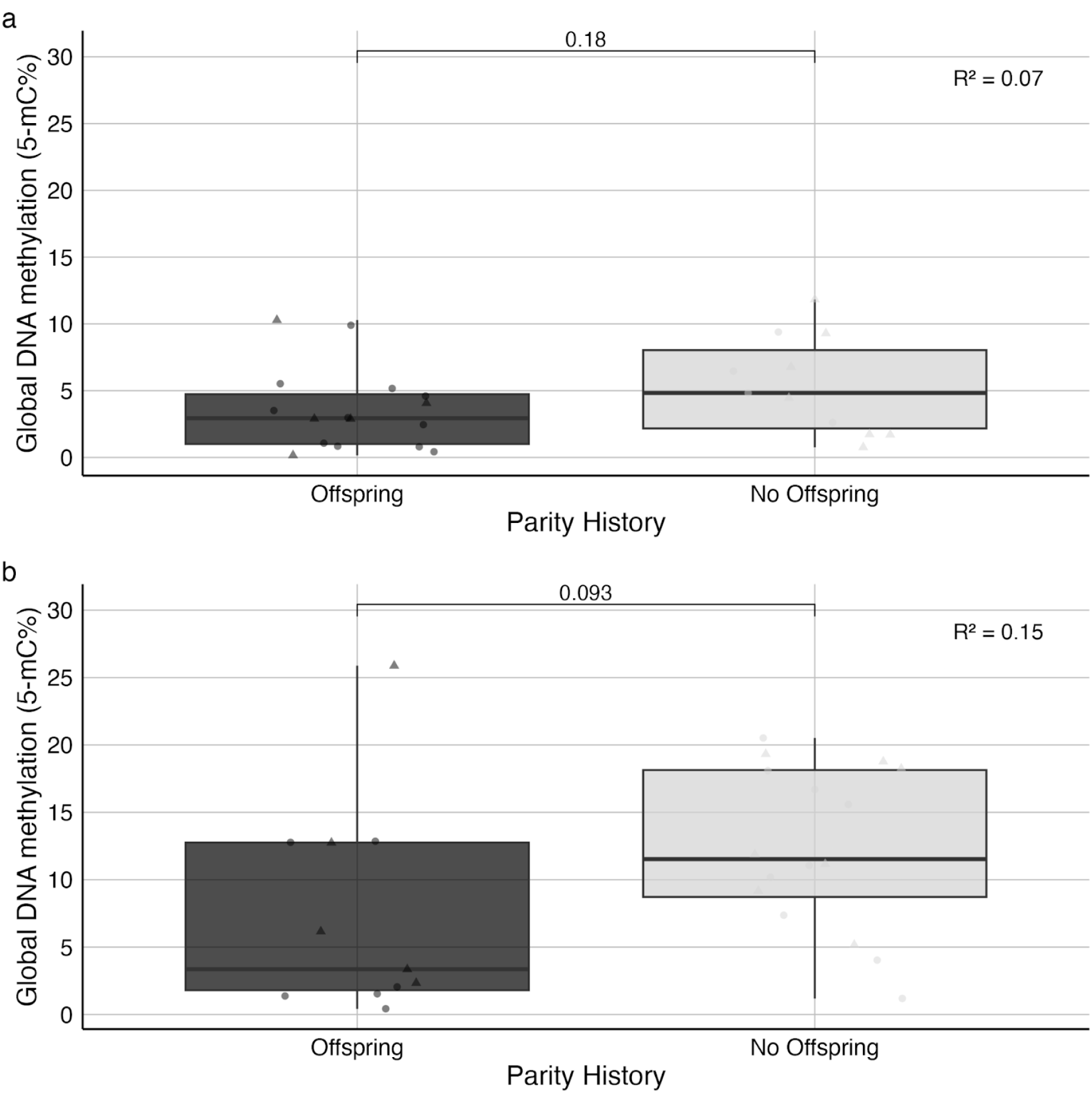
Differences between global DNA methylation in parents and non parents in blood and brain. a. Boxplot representing blood 5-mC% individuals with offspring (black boxplot n=20) and without offspring (grey boxplot, *n*=28). b) Boxplots representing brain 5-mC% in those with offspring (black boxplot n=11) and without offspring (grey boxplot, *n*= 19). For each box, the central mark represents the median; the lines delineate the 95% confidence interval of the median; edges indicate the 25th and 75th percentiles. Welch Two sample T-test demonstrated non-significant differences in mean/median in the brain ((t(17.71)=-1.78, p = 0.09)) and in blood (t(18.93.)=-1.38, p = 0.18)). Males are represented by circles and females are represented by triangles.

We then explored sex effects on 5-mC%. Both males and females parents showed lower content of global DNA methylation for both tissues, however, differences did not reach statistical significance (Blood males independent 2-sample T Test (t(16.90)=-0.97, p = 0.35), females with (M= 4.88, SD= 3.92) and without (M = 7.94, SD= 7.44) parity history (t(17.13)=-1.27, p = 0.22) or (brain: males (M= 5.17, SD= 5.95) and without (M = 11.63, SD= 6.61) parity history (t(11.67)=-1.97, p = 0.07), brain females with (M= 10.09, SD= 9.71) and without (M = 13.39, SD= 5.48) (t(5.83)=-0.69, p = 0.52). Due to small sample sizes, we could not distinguish between having one or several offspring on epigenetic ageing.

As the likelihood of being a parent increases with age, which is in turn a predictor of DNA hypomethylation, independent Welch Two Sample T-tests were used to test for differences in age distribution in parents and non-parents. These tests demonstrated a significant difference in the mean age between individuals with no offspring in blood (age_offspring_ = 13.15 years, SD = 5.59; age_no offspring_ = 7.79, SD = 4.04, t(32.61) = 3.6605, p < 0.05) and also in brain (age_parents_ =14.73, SD = 5.10); age_non parents_ = 6.53, SD = 2.22. t(12.79) = 5.28, p < 0.05).

#### Brain age related DNA methylation decline differs between parents and non-parents

We then used a linear regression to test the association between global DNA methylation decline and increasing ageing in both tissues (Figure 6a and 6b). As we did not identify differences between sexes, we combined ageing trajectories of females and males. A noticeable difference from scatter plots is age distribution for parents and non parents for both tissues (Fig. 6a and 6b). Furthermore, linear regression demonstrated that blood DNA methylation decline was negatively but non significantly associated with age for both parents and Non parents (blood non parents_5-mC%_ β=-0.75, 95% CI=[-1.71,0.20], p = 0.12, blood Parents_5-mC%_ (β=-0.29, 95% CI=[-0.67,0.095], p = 0.13).

**Figure 6.**
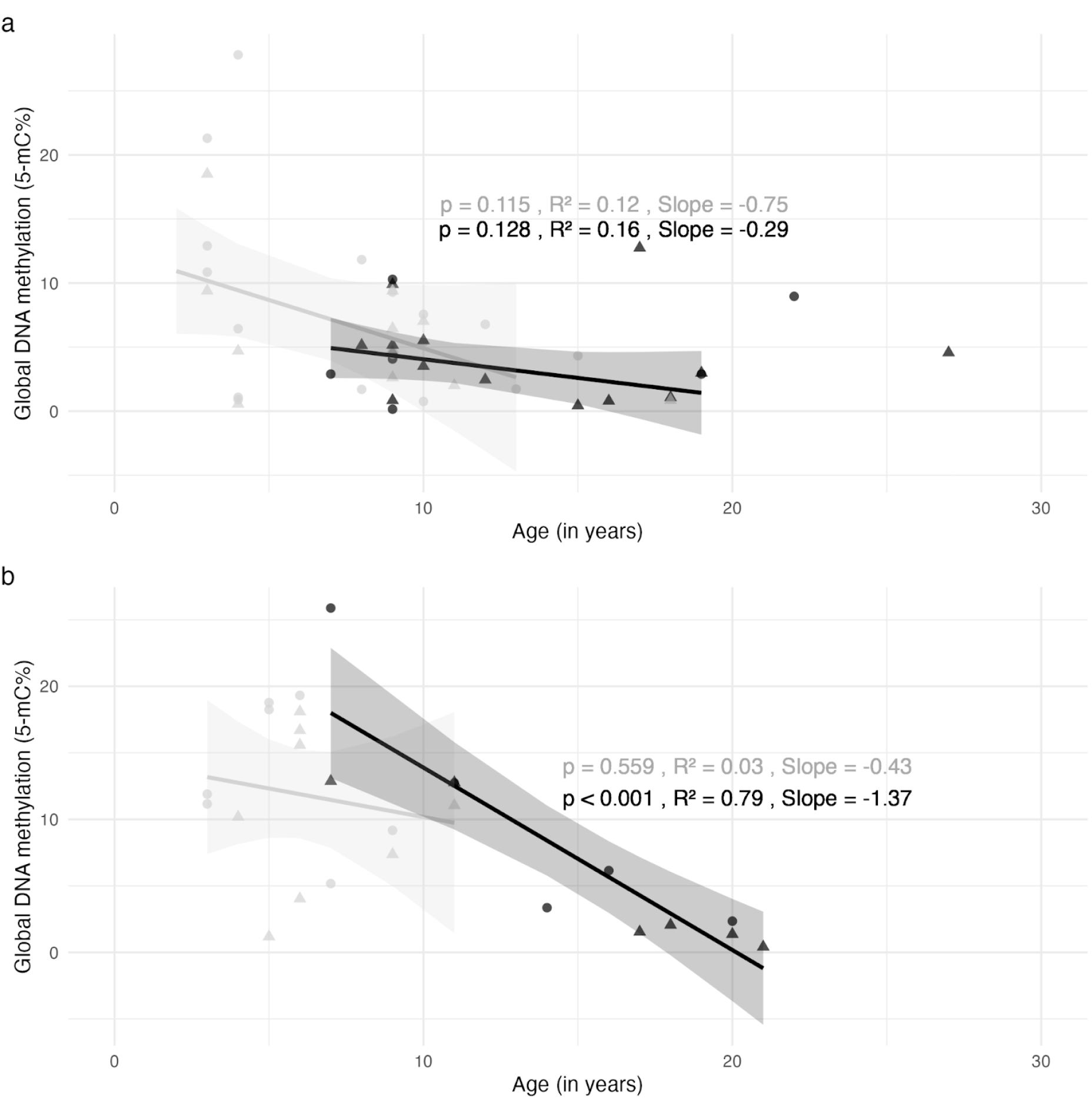
Global DNA methylation changes in blood and brain samples of different ages in parents and non parents. a) The relationship between global DNA methylation and age owl monkey blood samples from individuals with offspring (black regression line, *n*=20) and without offspring (grey regression line, (*n*=22). b) Brain global DNA methylation and age owl monkey brain samples from individuals with offspring (black regression line, *n*=11) and without (grey regression line, *n*=16). Males are represented by circle data points, females are represented by triangle data points. The size of the data points represent parental investment from none, small and high investment; small to large. Linear regressions demonstrated non-significant declines of blood 5-mC with age for both parents and non parents (p=0.12; p=0.13). For brain samples, there is a significant negative relationship between parents’ brains 5-mC% and increasing age (p value < 0.001) but this was not seen for non parents (p= 0.33).

We identified a contrasting pattern in the brain (Figure 6b). For non parents (grey regression line) 5-mC % was non significantly associated with age, and there was a mild decline (brain_non-parents_β=-0.43, 95% CI=[-1.98,1.12], p = 0.56), whereas for parents, the negative association between 5-mC% and age was significant (brain_parents_β=-1.37, 95% CI=[-1.91,-0.84], p < 0.001) with a steeper decline.

#### 4. Combination of ageing exposures does not accelerate epigenetic ageing in blood

Multiple regression models (2 and 3-way) were fitted to explore the cumulative effect of all the variables and interactions on epigenetic ageing. Analysis was conducted only in blood DNA methylation, for which we had data for three exposures (early life adversity, infections and parental investment). We predict that cumulative exposures would expedite the epigenetic ageing process.

In the three-way interaction model, the interaction between infection history, maternal rejection and parity on epigenetic ageing was examined. As seen with the individual linear regressions, age had a very close to significant hypomethylation effect on global 5-mC, suggesting a decrease in global 5-mC with increasing age. No significant effects were observed for sex, infection history, maternal rejection, or parity on global DNA methylation ageing trajectories. A non-significant positive association was found between having offspring and global DNAm indicating a potential increase in global 5-mC for individuals with offspring. An interaction effect was examined between infection history, maternal rejection, and parity. However, the combined interaction on epigenetic ageing did not reach statistical significance, suggesting that their joint effect does not significantly influence global 5-mC.

In the two-way interaction model, only the interaction between infection history and maternal rejection was examined. Similarly as seen in three way interaction, age did have a significant hypomethylation effect on global 5-mC (p<0.05). As shown in Table 1, there were no significant interactions with other factors.

**Table 1.** Three-way and two-way multiple regression models. For the three-way interaction model, we explored the combined influence of infection history, maternal rejection, and parity on epigenetic aging. In the two-way interaction model, we focused solely on the interaction between infection history and maternal rejection. Significance is indicated by asterisks (*), with *** (p < 0.001), ** (0.001 ≤ p < 0.01), * (0.01 ≤ p < 0.05), and . (0.05 ≤ p < 0.1). Three-Way Model: F = 1.21; df (8, 39); p = 0.32; R = 0.20; R-squared = 0.03. Two-Way Model: F = 1.64; df (6, 41); p = 0.16; R = 0.19; R-squared = 0.08.

| Variable | Coefficient<br>(Three-Way) | Std Error | t-value | p-value |
| --- | --- | --- | --- | --- |
| (Intercept) | 9.1675 | 3.1916 | 2.872 | 0.00663 ** |
| Age | -0.3699 | 0.1865 | -1.983 | 0.05460 . |
| Sex (Male) | -1.4972 | 1.7446 | -0.858 | 0.39616 |
| Infection History (Infected) | -0.7181 | 4.5337 | -0.158 | 0.87498 |
| Rejection Status (Rejected) | 3.8410 | 3.0656 | 1.253 | 0.21789 |
| Parity History | 0.4079 | 2.8441 | 0.143 | 0.88672 |
| Infection History:Rejection Status (Interact.) | -3.5160 | 5.7818 | -0.608 | 0.54673 |
| Infection History:Parity History (Interac.) | 3.8596 | 6.2352 | 0.619 | 0.53961 |
| Rejection Status:Parity History (Interaction) | -1.7458 | 3.7619 | -0.464 | 0.64524 |
| Infection History:Rejection Status:Parity History | -4.7635 | 9.2528 | -0.515 | 0.60966 |
| Variable | Coefficient<br>(Two-Way) | Std Error | t-value | p-value |
| (Intercept) | 9.30810 | 2.84091 | 3.276 | 0.00214 ** |
| Age | -0.35037 | 0.17251 | -2.031 | 0.04877 * |
| Sex (Male) | -1.52702 | 1.67779 | -0.910 | 0.36807 |
| Infection History (Infected) | 1.11160 | 3.04992 | 0.364 | 0.71738 |
| Rejection Status (Rejected) | 2.68441 | 1.78421 | 1.505 | 0.14011 |
| Parity History | -0.06888 | 1.97028 | -0.035 | 0.97228 |
| Infection History:Rejection Status (Interact.) | -4.96566 | 4.16705 | -1.192 | 0.24025 |

## 4. Discussion

In this study, we explored the potential of global DNA methylation measurements to act as surrogate markers of epigenomic ageing clocks, examining interaction of 5-mC% with ageing in blood and brain samples. Our findings confirm global DNA methylation declines with age in blood and in the brains of typical individuals. Furthermore, our study demonstrates that measurement of blood global DNA methylation responds to early life adversity, an example of environmental exposure, by accelerating its decline with age, and might underlie increased morbidity and mortality in this stressed cohort.

One of the key observations in our study was the confirmation of age-related declines in blood DNA methylation, consistent with previous reports in mammals. This decline, evident as a reduction of 0.33% in blood 5-mC content per year lived, reflects a widely observed phenomenon in ageing research. The Pearson correlation between chronological age and blood DNA methylation, explaining 10% of the variance, further emphasises the association between ageing and epigenetic changes. These results align with existing literature documenting age-related declines in global DNA methylation across various species (Maegawa et al., 2010; Hannum et al., 2013; Horvath et al., 2013; Paoli-Iseppi et al., 2017).

However, our study revealed an intriguing discrepancy between blood and brain tissue regarding DNA methylation dynamics with age. While blood exhibited the expected hypomethylation decline, consistent with previous findings, when studying all brain samples, we did not see the same declining trend. This unexpected result highlights the tissue-specific complexities of epigenetic regulation. Previous studies have reported overall reductions in global DNA methylation content in human brains (Heyn et al., 2012), but also differences in ageing patterns between cortical and pan-tissue epigenomic clocks (Grodstein et al., 2020; Shirevy et al., 2020) suggesting that brain ageing rates might differ to those seen in other tissues. Additionally, studies comparing brain and blood ageing patterns in humans and chimpanzees have suggested a slower rate of ageing in the brain relative to blood (Guevara et al., 2022), which might explain our results. It is also possible that life events experienced by owl monkeys sampled for our study might affect blood and brain differently, which would affect interpretation of these results.

Our exploration of the impact of maternal rejected owl monkeys on DNA methylation dynamics provided evidence of the value of measuring 5-mC% to detect epigenetic ageing acceleration linked with early life experiences. Consistent with the developmental origins of health and disease (DOHaD) framework, which proposes that early life conditions shape long-term health outcomes (Baker & Osmond, 1986) through epigenetic mechanisms (Gluckman and Hanson, 2011) our study demonstrated differences in DNA hypomethylation decline in blood samples from individuals exposed to maternal rejection (Figure 4b). In this owl monkey population, maternal rejection has been associated to increased mortality in infancy (Sanchez et al 2006), elevated cortisol secretion and frequency of treatments for infections (Osman et al 2021), and reduction in reproductive output, increased rates of maternal rejection, and early death in adulthood (Farinha 2023). Acceleration of global DNA methylation decline is then added to possible mechanisms which mediate increased morbidity and mortality in this cohort of rejected owl monkeys. Furthermore, our finding corroborates previous evidence suggesting a link between early life stress and accelerated epigenetic ageing in both humans and non-human animals (Bateson & Nettle, 2018, Pepper et al., 2018; Sheldon et al., 2022; Nettle et al., 2017). Considering the breadth of literature on the effect of early life adversity on behaviour, cognition and neuronal epigenome through the lifecourse (Maccari et at; 2014; Vaiserman & Koliada 2017) effects on brain epigenetic ageing of owl monkeys brains are likely, which will be assessed in future studies.

In contrast, our analysis did not reveal significant differences in DNA methylation dynamics between individuals treated for infections and healthy controls, although a trend towards accelerated epigenetic ageing was observed in the group treated for infections (Figure 4b). This finding underscores the complexity of environmental exposures and their differential effects on epigenetic regulation. Factors such as the type, duration, and severity of infections, as well as individual variability in immune responses, may contribute to the variability in DNA methylation patterns observed. Moreover, the small sample size of individuals treated for infections (n=9) and variability of treatments length and causing infecting agent not recorded adds to statistical noise, and limits the statistical power to detect significant effects, highlighting the need for larger studies with standardised protocols to elucidate the impact of infectious diseases on epigenetic ageing in this species.

Our next ageing exposure case study yielded striking results on the effects of parenting on ageing trajectories of different tissues. Individuals with a history of successful reproduction exhibit lower DNA methylation levels compared to non-parents, but this effect was only significant for brain tissue (Figure 5a and 5b). However, we also identified a significant difference in age distribution, with parents being significantly older than non parents, and this effect is seen for both blood and brain cohorts. As ageing is associated with DNA hypomethylation, we can not conclude from statistically lower DNA methylation in the brain of parents is attributable to parenting experience alone. We drew further insight from linear regressions analysis (Figure 6a and 6b). Starting our analysis in blood (6a) we observed an overall lack of overlap in age between parents and non parents in both tissues, which is expected for populations studied within the context of a breeding colony, where all animals that reach the age of sexual maturity are paired to produce young. Despite age differences, we in blood see the expected DNA methylation declining∼ageing trajectory, although this correlation was statistically insignificant for parents and non-parents, suggesting other factors contribute to blood global DNA methylation content. Inspection of linear regression of brain samples (6b) shows a markedly different picture. For non-parents there is a non-significant and mild decline (β=-0.43, R^2^=0.03) in global DNA methylation and age. However, measurement of 5-mC% in animals reaching parenting age is accompanied by a significant and strong decline (β=-1.37) in global DNA methylation content, with a strong association (R^2^=0.8) between increasing age and lower DNA methylation (Figure 6b). By comparing parenting effects across tissues we can conclude that significant ageing acceleration is induced by parenting in brains but not in blood. This result suggests that brain global DNA methylation does decline with age, when assessment is focused on ‘typical’ individuals that produce offspring in a breeding colony. Also our finding is consistent with the disposable soma theory of ageing, which proposes an evolutionary trade-off between reproduction and somatic maintenance (Kirkwood, 1977). Since in owl monkeys, parenting in males and females is associated with cognitive plasticity (Bardi et al., 2014), and in humans, parenting is associated with structural changes in prefrontal cortex in mothers (Kim et al., 2010), it is likely that parenting exerts greater ageing effects on the brain of owl monkeys when compared to blood.

Multiple regression analysis revealed significant DNA hypomethylation effects with age but did not detect significant acceleration effects of exposures on epigenetic ageing. The discrepancies between simple and multiple regression models could be explained by various factors. Including all variables in the one (multiple regression) model allowed us to examine their associations with global DNAm while controlling for the effects of each other. The lack of significant interactions between the variables could indicate that their combined effects are additive rather than interactive i.e. The presence of one stressor might have a similar effect on global DNAm regardless of the presence of other stressors. Additionally, compared to the simple/individual regressions, the multiple regression includes interaction terms that capture the combined effects of multiple variables and therefore the coefficients represent the change in global DNAm in one predictor/variable while holding the others constant. Then having more predictors/variables increased the model complexity compared to the simple regressions which captures more nuances in the data and so it is harder to interpret effects. As our sample size was small, it is likely that the multiple regression model did not have enough power to detect effects that were significant in the individual simple regressions (Jenkins and Quintana-Ascenio 2020; Serdar et al., 2021).

In conclusion, our results suggest global DNA methylation analysis provides a cost-effective approach to assess biological ageing, and captures the effect of ageing exposures in a tissue specific manner. This method requires only small blood samples and basic laboratory equipment available in diagnostic laboratories (elisa plate reader, DNA extraction), facilitating data collection from a greater number of individuals and populations.

Lastly, whereas this study shows how in an owl monkey colony DNA methylation trajectories respond to ageing exposures in captive settings, this knowledge can be extrapolated to primates facing stressors of the wild: habitat fragmentation (Arroyo-Rodriguez et al., 2010, Rangel-Negrin et al. 2014) and climate change (Kamilar & Beaudrot 2018) known stressors of primates. Testing how wild populations DNA methylomes age in response to ecological stressors warrants further investigation.

